# Three-dimensional cell neighbourhood impacts differentiation in the inner mass cells of the mouse blastocyst

**DOI:** 10.1101/159301

**Authors:** Sabine C. Fischer, Elena Corujo-Simón, Joaquín Lilao-Garzón, Ernst H. K. Stelzer, Silvia Muñoz-Descalzo

## Abstract

During mammalian blastocyst development, inner cell mass (ICM) cells differentiate into epiblast (Epi) or primitive endoderm (PrE). These two fates are characterised by the transcription factors NANOG and GATA6, respectively. Here, we present quantitative three-dimensional single cell-based neighbourhood analyses to investigate the spatial distribution of NANOG and GATA6 expression in the ICM of the mouse blastocyst. The cell neighbourhood is characterised by the expression levels of the fate markers in the surrounding cells, together with the number of surrounding cells and cell position. We find that cell neighbourhoods are established in early blastocysts and different for cells expressing different levels of NANOG and GATA6. Highest NANOG expressing cells occupy specific positions within the ICM and are surrounded by 9 neighbours, while GATA6 expressing cells cluster according to their GATA6 levels. The analysis of mutants reveals that NANOG local neighbourhood is regulated by GATA6.

**Summary statement:** Three-dimensional cell neighbourhood, which includes fate marker levels, number of neighbouring cells and cell position, determines cell fate decision in early mouse embryos.

## Introduction

During mammalian preimplantation development, two sequential cell fate decisions occur that result in three cell populations. Upon the first decision, cells become either trophectoderm (TE) or inner cell mass (ICM) cells. Descendants of TE cells form the foetal portion of the placenta, while the ICM cells make a further decision: they differentiate either into Epiblast (Epi) or into Primitive Endoderm (PrE). Epi cells predominantly give rise to the embryo proper while PrE cell descendants generate the endodermal part of the yolk sac (reviewed in ^1^). In mice, a three phase model has been proposed for ICM cell differentiation into Epi or PrE ^2, 3^. In early blastocysts, ICM cells co-express Epi and PrE markers such as NANOG and GATA6, respectively. Thereafter, in mid blastocysts, Epi progenitors express only NANOG, i.e. no GATA6, and PrE progenitors express only GATA6, i.e. not NANOG. During this phase, NANOG and GATA6 repress each other in a cell ^4-11^. Furthermore, FGF/MAPK signalling reinforces PrE commitment in PrE progenitors: Epi progenitors secrete FGF4, which binds to FGFR1 on Epi and FGFR2 on PrE biased cells ^12–15^. This results in the ‘salt-and-pepper’ distribution of Epi and PrE progenitors in the ICM without any obvious pattern ^2, 3^. Finally, in late blastocysts, the cells of the two lineages are segregated. PrE progenitors are polarised and positioned at the surface of the ICM, where they form an epithelium in contact with the blastocyst cavity or blastocoele ^16–18^. Epi versus PrE differentiation has been extensively studied in the context of marker expression dynamics and the involved signalling pathways (reviewed in ^19^). However, to our knowledge, the three-dimensional spatial distribution of NANOG and GATA6 expressing cells in the ICM has never been documented.

Technical developments have made it possible to study cell fate decisions during preimplantation mouse development at single-cell resolution (reviewed in ^20^). Invasive studies based on single cell transcriptomics have been used to investigate Epi versus PrE differentiation. However, transcriptomic techniques disrupt the cell positional information within the ICM ^12–14, 21^. A complementary approach is single cell resolution imaging based on immunofluorescence stainings ^2, 4, 6, 9, 14, 22–24^ or fluorescent reporters ^15, 25–28^. Combined with quantitative image analysis, the immunofluorescence approach provides protein expression levels and the positional information of a cell. So far, this type of data has not been analysed with respect to the cell neighbourhood, i. e. the expression level of the fate markers in the surrounding cells, the number of surrounding cells and cell position.

We recently proposed cell graphs as a tool to evaluate morphometric features of three-dimensional cell neighbourhoods in multicellular systems ^29^. Here, we extend this approach by including single cell expression levels of NANOG and GATA6. This enables the investigation of the three-dimensional spatial distribution of Epi and PrE progenitors in mouse embryos. To obtain a chronological order for data from fixed mouse embryos, we implemented a new method to follow cell fate specification based on the percentage of cells co-expressing NANOG and GATA6 in the ICM, which decreases as cells differentiate. In summary, our procedure allows the investigation of the spatial arrangement of Epi versus PrE differentiation of ICM cells as cell fate specification progresses.

To elucidate the ambiguous pattern of NANOG and GATA6 expressing cells in mid blastocysts, we simulated the three obvious models for random or alternating three-dimensional distributions of Epi and PrE progenitors and compared our simulations with the quantitative single cell data obtained from our experiments. This comparison revealed that none of the models is consistent with the experimental data at any stage. Instead, a more complex pattern of NANOG and GATA6 distribution in the cell neighbourhood is present in early blastocysts. High levels of NANOG expression in a cell correlate with a specific number of neighbours and location within the ICM. The GATA6 level of any given cell correlates with the levels of GATA6 in its neighbours, resulting in GATA6 expressing clusters of cells. Finally, through the use of mutants, we show that the NANOG neighbourhood is regulated by GATA6.

Altogether, we present Epi versus PrE differentiation in relation to cell position in the ICM, relative to nearby cells and their fate markers expression levels. Our results highlight GATA6-dependent mechanisms for the spatial arrangement of NANOG and GATA6 expression in the ICM. The defined spatial arrangement is already present in the early blastocysts and allow the cells to make a coordinated fate decision depending on the number of neighbours and their expression levels, together will cell position.

## Results

### Quantitative three-dimensional analyses of spatio-temporal patterns of NANOG and GATA6 during mouse preimplantation development

In this study, we developed a pipeline for the quantification of the three-dimensional spatial distribution of cell fate markers, taking into account the single cell level as well as the cell neighbourhood (Fig. 1A). The quantitative immunofluorescence (QIF) analysis of NANOG and GATA6 at the single cell level in mouse preimplantation embryos in different stages of development using MINS provided the cell positions and the expression levels (Fig. 1A (I-II)-B, ^30^). This information was implemented into a Delaunay cell graph for each embryo, in which vertices represent cells and edges represent neighbourhood relations between cells (Fig. 1A (III-IV), ^29^). Implementing a Delaunay cell graph allows an approximation of which cells (nuclei in this case) are in physical contact ^29, 31, 32^.

**Fig. 1.**
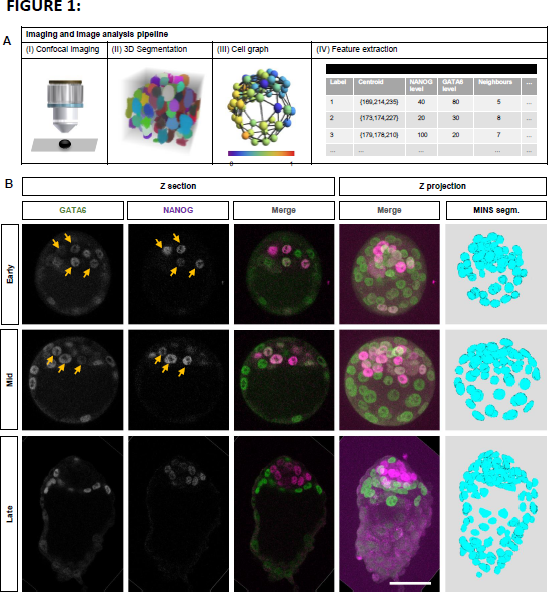
Three-dimensional imaging-based quantitative cell neighbourhood analysis of Epiblast vs Primitive Endoderm fate differentiation in preimplantation mouse embryos. **(A)** Image analysis pipeline for multiscale characterisation of mouse embryos. (I) Whole embryos were imaged using a confocal laser scanning fluorescence microscope using sequential scan mode. (II) Embryo images were segmented with MINS ^30^. (III) Graphical representation of the Delaunay cell graph for one of the embryos. (IV) Feature extraction of each individual cell of each embryo including the centroid position of each nucleus and markers’ intensity values. **(B)** Representative confocal images of mouse preimplantation embryos immunostained for GATA6 (green, PrE marker) and NANOG (magenta, Epi marker) at the indicated developmental stages. Yellow arrows point to cell co-expressing NANOG and GATA6. The first three columns are single confocal z-sections, the last columns show the maximum z-projection of the merged confocal and the segmented images using MINS on the DAPI channel (not shown). Scale bar: 50µm. These embryos were all imaged in the same confocal session and processed exactly the same way.

We used a total of 45 embryos from three independent experimental replicas, imaged in four confocal sessions (Fig. 2A). Given the observed quantitative differences between replicas due to variability in the experimental and imaging setup, we aligned the data according to NANOG and GATA6 threshold values for each experiment (Fig. 2B, see Materials and Methods). Based on the common thresholds, we identified double positive (DP: N+ and G6+), double negative (DN: N- and G6-), NANOG+/GATA6-(N+/G6-) and NANOG-/GATA6+ (N+/G6-) cell populations.

**Fig. 2.**
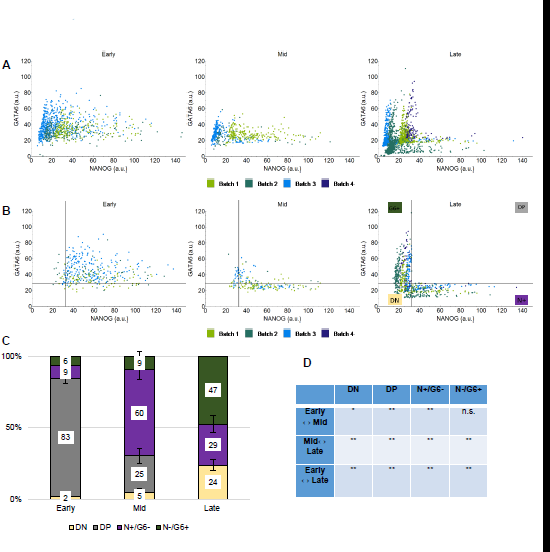
Quantitative population analysis of ICM cells in preimplantation mouse embryos. **(A)** Scatter plots showing NANOG (x-axis) and GATA6 (y-axis) levels in all cells in early, mid and late blastocysts (left, centre and right, respectively) in arbitrary units (a.u., here and in subsequent similar graphs). Each dot represents the levels in a single cell from 19 early, 10 mid and 16 late blastocysts. **(B)** Scatter plots showing NANOG (x-axis) and GATA6 (y-axis) levels in ICM cells in early, mid and late blastocysts (left, centre and right, respectively) after aligning the data sets. Dashed lines represent the threshold levels for NANOG and GATA6 calculated by a cluster analysis on the late blastocyst data (see Materials and Methods). The data were pooled from the experiments shown in A. **(C)** Proportion of ICM cells expressing GATA6 (N-/G6+), NANOG (N+/G6-), co-expressing NANOG and GATA6 (double positive, DP) or neither (double negative, DN) in ICM cells in early, mid and late blastocysts. Staging was performed according to the percentage of DP cells (see Materials and Methods). Error bars correspond to the standard error of the mean here and in subsequent similar graphs. See also Fig. S1 and S2. **(D)** Statistical analysis comparing the proportion of cells in the different populations in the indicated stages using a z-test with Bonferroni correction; **: p<0.01; *: p*0.05; n.s.: not significant.

The next step was to group embryos in a temporal sequence to investigate Epi versus PrE differentiation as a function of time. In the past, two different staging methods have been proposed. Mouse preimplantation embryos can be staged by days after fertilization (E0.5-E4.5) ^33^ or the total number of cells^3^. However, the actual time point of the fertilization is not determinable, and does not take into account the developmental variation between embryos from the same female ^3^ or from different strains ^34^. Staging our data set using total cell number regardless of the proposed cell ranges used ^3, 6, 13, 24, 27^ resulted in no decrease in the percentage of DP cells from early to mid blastocysts as we recently proposed (not shown, and ^24^. Furthermore, we noticed that in several embryos that would be classified as mid blastocysts using either staging method, almost all ICM cells were DP in this data set and previous data sets (not shown and ^24^).

Therefore, we used the cellular event staging method to stage the embryos based on Epi versus PrE specification progression, i. e. decrease in the DP population. Early blastocysts (average total cell number 49±2.5 cells) have a majority, hence more than 55% DP cells in the ICM, and mid blastocysts (50±5.7 cells) have less than 55% DP cells in the ICM. Late blastocysts (139±6.8 cells) have no DP cells in the ICM and PrE cells are aligned next to the blastocoele (cavity). Using this staging method, we split our 45 data sets into 19 early, 10 mid and 16 late blastocysts. During the transition from early to mid blastocyst stage, the proportion of DP cells decreased significantly and the proportion of N+/G6- and N-/G6+ cells increased significantly (Fig2C-D and Fig. S1), as expected ^24^. We also checked how NANOG and GATA6 levels changed as cell fate specification progresses: NANOG decreased while GATA6 did not (Fig. S2). These results fit with the previously proposed mutual inhibition of NANOG and GATA6 in each cell ^4, 5, 7-9, 11, 35^. Assuming such a model, initial conditions with positive NANOG and GATA6 can result in steady states with similar or reduced levels of NANOG or GATA6, respectively (Fig. S3).

In summary, we implemented an unbiased three-dimensional data analysis pipeline for mouse blastocysts that integrates QIF measurements with the spatial arrangement of three-dimensional cell neighbourhoods. We also suggested a new staging method based on the progression of ICM cells towards Epi or PrE fate. Admittedly, this method is not applicable for *Nanog* or *Gata6* mutant embryos. Nevertheless, it presents a series of advantages for the wild type. Unlike the cell number staging method, this cellular event staging method can be used in aggregation chimeras ^36^, as well as in embryo splitting experiments ^37^. Furthermore, this method allows a more direct comparison between different species with different embryonic times or sizes that also show co-expression of NANOG and GATA6, such as marsupials ^38^, cows ^39^, marmosets ^40^, and even humans ^41^.

### Three-dimensional cell neighbourhood analyses challenge the salt-and-pepper distribution model of NANOG and GATA6 expressing cells in the ICM

A salt-and-pepper distribution of N+/G6- and N-/G6+ cells has been proposed as a description of their distribution in mid blastocyst embryos ^2^. How this salt-and-pepper distribution develops in two dimensions ^9^ and resolves during cell sorting in three dimensions has been modelled ^42^. However, a precise definition has never been presented.

Based on the prevailing notions of a salt-and-pepper pattern, we simulated three types of reference patterns. We considered only two types of expression states: positive or negative. These are assigned to ICM cells according to the following rules, taking into account their three-dimensional positions obtained above (see Materials and Methods, Fig 3A):

– A random model, in which positive and negative expression states are randomly distributed in the ICM (Fig. 3A, left panel).
– A period two model ^43^), in which cells with positive expression state are surrounded by cells with negative expression state (Fig. 3A, central panel).
– A nearest neighbour model, in which one cell is assigned a state and its nearest neighbour from the remaining cells without state is assigned the opposite state, until there are no cells left (Fig 3A, right panel).

**Fig. 3.**
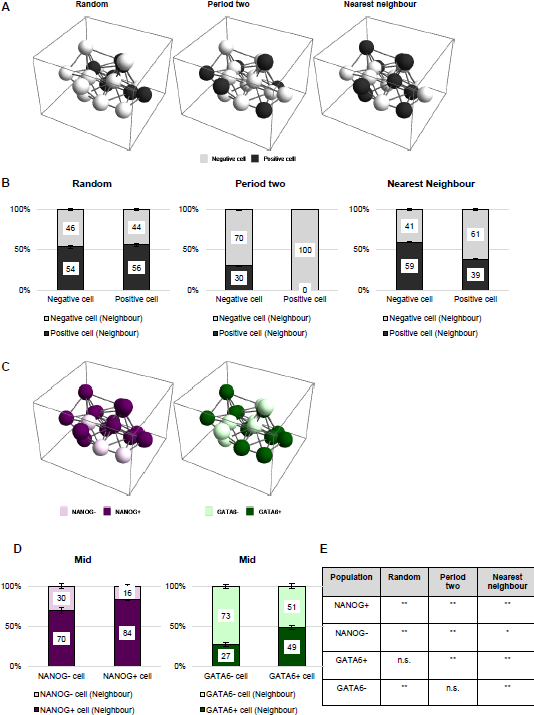
N+/G6- and N-/G6+ cell distribution in ICMs in mid blastocysts does not show a salt-and-pepper pattern. **(A)** Three-dimensional representation of simulated patterns of ICMs. The patterns generated by using the cell positions of a representative mid blastocyst embryo from the data and one of the reference patterns: a random distribution of positive and negative cells (left), a period two distribution (centre) and a nearest neighbour distribution (right, see text for details). Black spheres represent positive cells and white spheres represent negative cells. **(B)** Proportion of positive or negative neighbours of positive or negative cells in the simulated patterns indicated in A. The numbers are based on simulated patterns for all mid blastocysts data sets from our embryo data. For the random simulation, GATA6 distribution is shown for simplicity (the simulation using NANOG is equivalent, not shown). **(C)** Three-dimensional representation of N+/N-cells (left) and G6+/G6-cells (right) of the ICM of a representative mid blastocyst embryo. The positions of the cells are the same as in A. In the left panel, N+/G6- and DP are considered as N+ (dark magenta) and N-/G6+ and DN as N-(light magenta). In the right panel, N-/G6+ and DP are considered as G6+ (dark green) and N+/G6- and DN as G6-(light green). These cell states are assigned according to the embryo data. **(D)** Proportion of N+ or N-neighbours of N+ or N-cells (left) and G6+ or G6-neighbours of G6+ or G6-cells in the ICM (right) of mid blastocyst embryos. See also Fig. S4. **(E)** Summary results of the Mann-Whitney statistical tests with Bonferroni correction comparing the simulated and experimental data proportions obtained in the mid blastocysts. **: p<0.01; *: p<0.05; n.s.: not significant.

Based on our Delaunay cell graph for each embryo, we calculated for each reference pattern the percentage of positive and negative neighbours of positive and negative cells by applying standard methods of spatial statistics (Fig. 3B, ^44^). We were interested in the spatial distribution of NANOG and GATA6 in the ICM, therefore, we only considered ICM cells for the cell of interest and its neighbours. For a direct comparison between the simulated and experimental data, we split the *in vivo* cells into N+ and N-as well as G6+ and G6-(Fig. 3C-D). N-cells include DN cells and N-/G6+ cells, while N+ cells are DP and N+/G6-cells. For G6+ and G6-cells, the assignment is analogous, i.e. DN and N+/G6-cells are G6-cells, and DP together with N-/G6+ are G6+ cells. The statistical comparison of the distribution of N+ and N-cells in *in vivo* ICMs with the distributions in the simulations shows that they differ significantly from any of the simulated patterns. The equivalent comparison for G6+ cells indicates that these cells are distributed following a random pattern, while G6-cells follow a period two pattern. In early blastocysts, we observe significant differences in neighbour distributions between embryo and simulated data in all cases (Fig. S4).

Our comparison between the spatial distributions of cell fate markers in the ICM and simulated patterns revealed that in the mid blastocysts, N+ and N-cells were not organized following a salt-and-pepper pattern. G6+ and G6-cells followed a combination of the random and the period two pattern. Epi and PrE progenitors displayed different three-dimensional spatial arrangements in the ICM, which suggests that their distribution might be regulated independently.

### Higher levels of NANOG in a cell do not instruct its direct neighbours into expressing GATA6

A salt-and-pepper pattern for Epi and PrE progenitors could not be detected in the ICM of early and mid blastocysts. Therefore, we next investigated the complete local neighbourhood of a cell, as the behaviour of an ICM cell is influenced by all its neighbours. Hence, for all following analyses, we considered cells from the ICM and all their neighbours, i.e. both ICM cells and TE cells.

It has been proposed that Epi fate reinforces PrE fate in neighbouring cells via FGF4 in mid blastocysts ^4, 22, 45, 46^. NANOG expressing cells would secrete FGF4, inducing GATA6 expression in the neighbouring cells. This hypothesis would predict a strong correlation of the levels of NANOG of a cell with the levels of GATA6 in its neighbouring cells. We used Spearman’s correlation coefficient, which measures monotonic relationships between two variables, to analyse the levels of NANOG and GATA6 relative to the respective levels in the neighbouring cells ^47^. The correlation analyses of GATA6 levels in ICM cells and median NANOG in its neighbours (regardless of these being TE or ICM) and *vice versa* are either very weak or weak in all cell population types and developmental stages (Fig. 4A-B, see Material and Methods for the classification of correlation strengths).

**Fig. 4.**
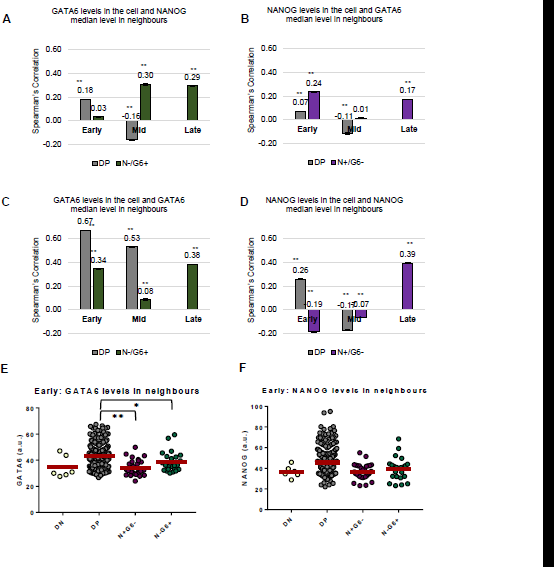
ICM cells in early blastocysts cluster together according to their GATA6 levels. **(A-B)** Spearman’s correlation coefficients of GATA6 (A) or NANOG (B) levels of a cell and the median NANOG (A) or GATA6 (B) levels of its neighbours in the indicated populations in the ICM at different embryonic developmental stages. The error bars represent the standard error calculated by bootstrap sampling 10.000 times of the experimental data, here and in subsequent similar graphs. **: p<0.01, Mann-Whitney test with Bonferroni correction between the experimental data and a random model, here and in subsequent similar graphs. **(C-D)** Spearman’s correlation coefficients of GATA6 (C) or NANOG (D) levels of a cell and the median GATA6 (C) or NANOG (D) levels of its neighbours in the indicated populations in the ICM at different embryonic developmental stages. **(E-F)** Scatter dot plots showing the expression levels of GATA6 (E) and NANOG (F) in neighbours of cells for the four indicated populations in early blastocysts. The red lines indicate the mean, here and in subsequent similar graphs. Mann-Whitney test with Bonferroni correction between the different populations: **: p<0.01; *: p<0.05. For expression levels of neighbours in other stages see Fig. S5.

These results were not consistent with the previously proposed hypothesis that Epi progenitors induce PrE fate in their direct neighbours at any preimplantation stage. Furthermore, they implied that although NANOG expressing cells are the primary source of PrE inducing signal, this signal does not activate the PrE programme in the direct neighbours of NANOG+ cells.

### ICM cells in early blastocysts cluster together according to their GATA6 levels

Since correlations for opposite fates could not be detected in the local cell three-dimensional neighbourhood, we next investigated correlations for the same fate ^6^. For this, we performed Spearman’s correlation analyses of the levels of NANOG or GATA6 in an ICM cell with the median NANOG or GATA6 levels of its neighbours (both ICM and TE neighbours), respectively (Fig. 4C-D). The correlation analysis of the GATA6 level in a cell and the median GATA6 level of its neighbours indicated moderate and strong positive correlations (Fig. 4C). The strong positive correlation was present in the DP population of early blastocysts and decreased to a moderate positive correlation in mid blastocysts. To ensure that any correlation that we detect is significant, we compared it to a null model with random expression levels (see Materials and Methods). This is the first time that local correlations for cells according to their GATA6 levels have been documented for early blastocysts. Repeating the analysis without the TE to rule out an effect of GATA6 expression in those cells (Freyer et al., 2015; Koutsourakis et al., 1999; Plusa et al., 2008) resulted in similar correlation results (data not shown). For NANOG, we detected only very weak or weak correlations and anti-correlations for all populations and stages (Fig 4D). In the late blastocysts, we observed only weak local correlations of NANOG or GATA6 levels. This can be explained by the fact that, at this stage the PrE (G6+) cells form an epithelium facing the blastocoele on one side and Epi (N+) cells on the other, while Epi cells start downregulating NANOG expression.

To further characterise the local cell neighbourhood based on their GATA6 expression in early blastocysts, we measured the levels of GATA6 in the neighbours of the four cell populations present in the ICM (Fig. 4E and Fig. S5). In early blastocysts, we observed that G6+ cells have neighbours with higher levels of GATA6, while G6-cells are surrounded by cells with low levels of GATA6. In mid and late blastocysts, this difference is statistically significant (Fig. S5A). The equivalent analysis for NANOG levels shows no clear trend in early blastocysts (Fig. 4F, and Fig. S5B).

In summary, our results showed that GATA6 expressing cells (both DP and N-/G6+) in the ICM were spatially organised according to their GATA6 expressing levels, while the same was not true when looking at NANOG. The local arrangement of G6+ cells was already present in the early blastocysts. It was characterized by the correlation of GATA6 levels between neighbouring cells indicating that cells with similar GATA6 levels clustered together. Our analyses do not distinguish whether this clustering is due to cell-cell communication, active or passive sorting, related lineage, or a combination of these. However, they provide a framework for investigating the contribution of these processes in early embryos to Epi versus PrE differentiation.

### Relationship between NANOG expression levels and relative cellular position within the ICM

We next tested whether mechanical and/or positional cues emerging from the ICM govern NANOG and GATA6 expression levels. For this, we analysed the relation between expression levels and total number of neighbours, and the distance to the ICM centroid (Fig. 5). We observed, particularly in early blastocysts, maximal NANOG expression in cells with 7 to 14 neighbours, with the highest level located in cells with 9 neighbours, independently of which markers levels these neighbours expressed (Fig. 5A). Visualisation suggested that the highest NANOG expressing cell in each embryo is located away from the ICM centroid in early blastocysts and closer to in mid blastocysts (Fig. S6). The detailed positional analysis of all ICM cells confirmed that in early blastocysts cells located around 20μm away from the ICM centroid expressed higher levels of NANOG (Fig. 5B). In mid and late blastocysts the highest NANOG expressing cells are located closest to the ICM centroid. Performing the same analyses for GATA6 expression levels, we detected no clear dependence between its expression levels and the total number of neighbours at any stage (Fig. 5C). The position of the highest expressing GATA6 changed as development progresses: no clear positional effect was observed in early blastocysts while in mid and late blastocysts, the highest GATA6 expressing cells are located furthest away from the ICM centroid, consistent with the sorting process (Fig. 5D).

**Fig. 5.**
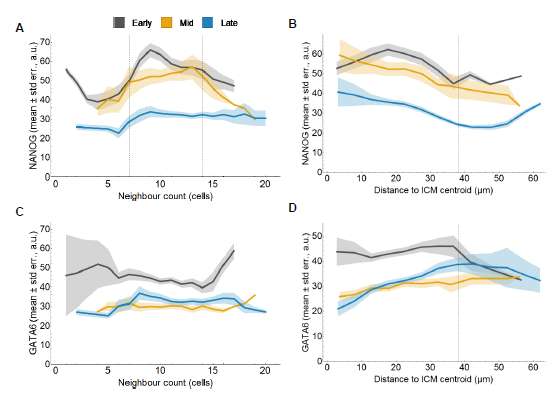
Relationship between NANOG expression levels and relative cellular position within the ICM. **(A,C)** Mean level of NANOG (A) or GATA6 (C) (y-axis) versus the number of neighbours (x-axis) for ICM cells in early, mid and late blastocysts. The shaded regions display the standard error of the mean, here and in subsequent similar graphs. **(B,D)** Mean level of NANOG (B) or GATA6 (D) (y-axis) versus the distance to the centre of the ICM (x-axis) for ICM cells in early, mid and late blastocysts.

In summary, these results showed that there is a clear interrelation between cell position within the ICM and NANOG expression levels, which changed as development progresses: away from the centroid in early and near the centroid in mid blastocysts. These changes in localization might be due to cell migration and/or upregulation/downregulation of NANOG in specifically located cells as development progresses. Altogether, this indicated that positional information regulates NANOG expression or that NANOG expression levels determined cell position in the ICM and hence Epi fate.

### Three-dimensional analyses with larger data set confirm the three-dimensional pattern of ICM cells

To ensure the robustness of our observations, we reanalysed a larger data set generated in a different laboratory with slightly different experimental set up (Fig. 6 and Fig. S7–S11, ^24^). Staging the embryos by progression of Epi versus PrE fate resulted in 70 early, 53 mid and 24 late blastocysts. As cell fate specification progresses, the percentage N-/G6+ and DN increased (Fig. S7A).

The three-dimensional neighbourhood analysis of GATA6 expressing cells confirmed our previous results with the first data set: the cells are clustered according to their GATA6 expression levels in early embryos (Fig. 6A-B). In the correlation analyses, we observed moderate and strong positive correlations of GATA6 levels in a cell with GATA6 levels of neighbouring cells in the DP cells and in the N-/G+ cells (Fig 6A). The analyses of GATA6 levels in the neighbours of the four cell populations showed that G6+ cells are surrounded by cells expressing higher levels of GATA6 (Fig. 6B, see Fig. S7B for other stages). Unlike with the first data set, here we observed very weak or weak correlations in mid blastocysts. We cannot rule out that this difference might be due to the differences in the data set size. The equivalent analyses with NANOG expressing cells showed again very weak or weak correlations and no clustering effect in early blastocysts (Fig. S7C-D).

**Fig. 6.**
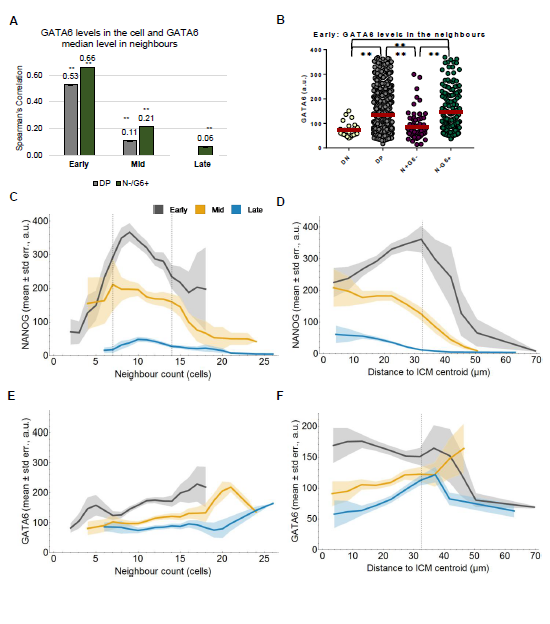
Three-dimensional neighbourhood analyses of data from Saiz et al. 2016 ^24^ data set. **(A)** Spearman’s correlation coefficients of GATA6 levels of a cell and the median GATA6 levels of its neighbours in the indicated populations in the ICM at different developmental stages. **: p<0.01, Mann-Whitney test with Bonferroni correction. See also Fig. S7C. **(B)** Scatter dot plots showing the expression levels of GATA6 in neighbours of cells for the four indicated populations in early blastocysts. Mann-Whitney test with Bonferroni correction between the different populations: **: p<0.01; *: p<0.05. For GATA6 and NANOG expression levels of neighbours in other stages see Fig. S7. **(C,E)** Mean level of NANOG (C) or GATA6 (E) (y-axis) versus the number of neighbours (x-axis) for ICM cells in early, mid and late blastocysts. **(D,F)** Mean level of NANOG (F) or GATA6 (H) (y-axis) versus the distance to the centre of the ICM (x-axis) for ICM cells in early, mid and late blastocysts.

The three-dimensional neighbourhood analysis of NANOG expressing cells reinforced the conclusion that NANOG expression levels related to cell position within the ICM (Fig. 6C-D, and Fig. S9). Again, we observed that already in early blastocysts, cells expressing highest NANOG levels had around 9 neighbours in early and mid blastocysts (Fig. 6C). Furthermore, these cells were away from the ICM centroid in early blastocysts and close to the centroid in the mid blastocysts (Fig. 6D). The equivalent analyses taking into account GATA6 expression levels showed again only a positional effect in mid and late embryos, where cells furthest away from the centre of the ICM have high levels of GATA6 (Fig. 6E-F). See also Fig. S10 for three-dimensional representation of selected embryos showing N+ and N- or G6+ and G6-expressing cells and their position in each developmental stage.

The statistical comparisons with simulated reference patterns of the prevailing notions of a salt-and-pepper pattern (Fig. 3A) showed again that the N+ and N-did not exhibit a salt-and-pepper pattern neither in early nor in mid blastocysts (Fig. S8A-F). The equivalent comparison for G6+ cells indicated that these did not follow any salt-and-pepper pattern either.

Testing whether Epi progenitors induced PrE fate in their direct neighbours at any preimplantation stage, showed again negative results (Fig. S8G-H). The correlation analyses of GATA6 levels in ICM cells and median NANOG in its neighbours (regardless of these being TE or ICM) and *vice versa* were either very weak or weak in all cell population types and developmental stages.

In summary, the three-dimensional neighbourhood analyses of a larger, independent data set validated our previous observations: 1) local coordination of NANOG and GATA6 expression is already present in early blastocysts, but with different features for Epi and PrE progenitors; 2) PrE progenitors form local clusters; 3) Epi progenitors are localized away from the centre of the ICM; 4) ICM cells in mid blastocysts show no salt-and-pepper pattern; 5) Epi progenitors do not induce PrE fate in their direct neighbours at any preimplantation stage.

### GATA6 regulates NANOG cell neighbourhood

The larger data set allowed us to analyse if the observed pattern in the location of NANOG expressing cells was specific to a cell population (Fig. S11). The peak of NANOG expression depending on total neighbours number is clearly defined both in the DP and in the N+/G6-population of early blastocysts (Fig.S11A-B). In the mid blastocysts, cells with more than 14 neighbours show lower levels in NANOG expression upon GATA6 expression (DP population), but not in its absence (N+/G6-cell population). This suggests that GATA6 might be needed for the downregulation of NANOG in cells surrounded by a large number of neighbours. To investigate this further, we analysed the neighbourhood of NANOG expressing cells in the absence of *Gata6* using the previously published data set composed by 20 *Gata6^+/+^*, 28 *Gata6^+/-^*, and 15 *Gata6^-/-^* embryos ^6^.

The most striking result is obtained when analysing NANOG correlations (Fig. 7A): the usual weak correlation found in early *Gata6^+/+^* blastocysts becomes a strong correlation upon reducing GATA6 levels to half (*Gata6^+/-^*), and remains strong in the complete absence of GATA6 (*Gata6^-/-^*) early blastocysts. In mid and late blastocysts this behaviour is maintained: the correlations are increased upon reducing GATA6 levels, although in the late ones the complete absence of GATA6 is needed in order to see an effect. These results suggest that GATA6 inhibits the coordination of NANOG levels in the neighbourhood. In other words, the spatial NANOG heterogeneity is lost upon decreasing or eliminating GATA6 and, cells cluster according to their NANOG levels.

**Fig. 7:**
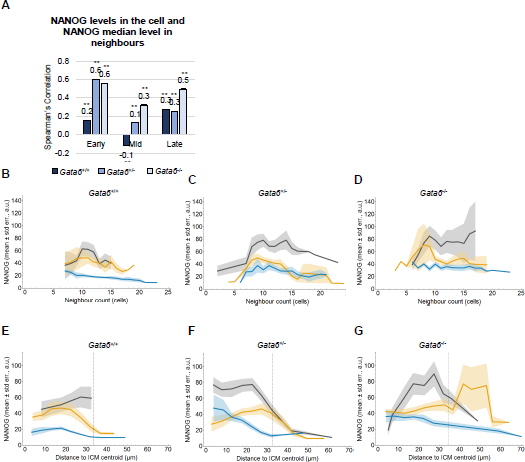
GATA6 regulates NANOG neighbourhood patterning: ^6^ data set. **(A)** Spearman’s correlation coefficients of NANOG level of a cell and the median NANOG levels of its neighbours in N+ populations (DP and N+/G6-) in the ICM at different embryonic developmental stages in *Gata6^+/+^*, *Gata6^+/-^* and *Gata6^-/-^* embryos. **: p<0.01, Mann-Whitney test with Bonferroni correction. Note that in some cases the error bars are so small that they cannot be appreciated in the figure. **(B-D)** Mean level of NANOG (y-axis) versus the number of neighbours (x-axis) for ICM and Epi cells in in *Gata6^+/+^* (B), *Gata6^+/-^* (C) and *Gata6’^-/-^* (D) early, mid and late blastocysts. **(E,G)** Mean level of NANOG (y-axis) versus the distance to the centre of the ICM (x-axis) for ICM cells in *Gata6^+/+^* (B), *Gata6^+/-^* (C) and *Gata6’^-/-^* (D) early, mid and late blastocysts.

The trivial explanation for this would be that upon reducing GATA6 levels more cells become NANOG+. However this was not the case, as the population analyses showed that the proportion of DN increases upon reduction of GATA6 and the proportion of NANOG+ cells stays similar (Fig. S11E). These DN cells often express CDX2 as a result of delayed downregulation ^6^.

The NANOG levels analysis versus the numbers of neighbouring cells also showed surprising results, especially in the early blastocysts (Fig. 7B-D): the reduction of GATA6 levels affects the relation between NANOG expression levels and the number of neighbours. The peak of highest NANOG expression around 9 neighbours gets wider in heterozygous early blastocysts (Fig. 7B-C). In mutant early blastocysts, unlike in the previous situations, cells with more than 14 neighbours maintain high NANOG expression (Fig. 7D). The position of these high NANOG expressing cells within the ICM in the early mutant embryos is still away from the centroid as in the *Gata6^+/+^* and *Gata6+/-* (Fig 7E-G). However in the mid blastocysts, they are localized even further away instead of closer to the centroid.

Altogether, these results suggested that GATA6 was involved in regulating NANOG neighbourhood pattern. The regulation was at multiple levels: 1) inhibiting the coordination of NANOG levels in neighbouring cells and, hence, its spatial heterogeneity; 2) downregulating the levels of NANOG in cells with a large number of neighbours; and 3) coordinating NANOG levels and cell position within the ICM.

## Discussion

Previous publications investigating Epi versus PrE fate decision in mouse embryos have focused on an imaged-based single cell immunofluorescence quantification of NANOG and GATA6 expression levels and/or the correlation between them (^3, 6, 9, 14, 24, 30^, and reviewed in ^20^). Here, we extended the single cell quantification to include three-dimensional neighbourhood analyses to evaluate the global spatial distribution of NANOG and/or GATA6 ^29^. Ultimately, our novel three-dimensional analyses allow us to propose a model of how Epi and PrE fates arise from the early blastocyst based on cell neighbourhoods and relative cell position (Fig. 8).

**Fig. 8.**
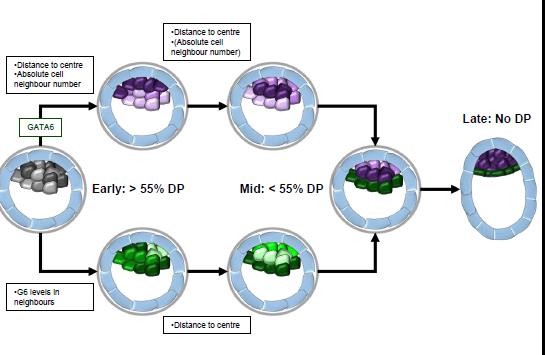
Three-dimensional spatial arrangement of NANOG and GATA6 expressing cells during cell fate decision in the ICM from early to late blastocysts. In early blastocysts, the majority (>55%) of the ICM co-express NANOG and GATA6 but at different levels. NANOG levels in a cell will depend on the distance to the centroid and the absolute cell neighbour number: cells away from the centroid and with 9 neighbour will express higher NANOG levels. These NANOG expressing cells’ pattern is regulated by GATA6. GATA6 levels in a cell will depend on the levels expressed in its neighbours ensuring that cell with similar levels cluster together. In mid blastocysts (<55% of DP cells), the highest NANOG expressing cells are located closest to the centroid, while GATA6 expressing cells will be already facing the blastocoele. These pattern are stablished in early and evolving in mid blastocysts and will be resolved in late blastocysts in preparation for implantation. See also Fig. S10.

### Spatial pattern of NANOG and GATA6 expressing cells in the ICM of early blastocysts

Our novel single cell quantitative three-dimensional neighbourhood analyses revealed a pattern in the ICM cells based on NANOG and GATA6 expression in early blastocysts (Fig. 8). Although most of ICM cells co-express both markers, the levels of each one vary among the different cells, as well as how these levels are regulated.

The spatial pattern of cells depending on their NANOG expression levels correlate with cell position. In early embryos, cells away from the centroid and with 9 neighbouring cells (regardless of their identity) display highest NANOG levels in early embryos (Fig. 8). There are several mechanisms that could link the positional information to NANOG expression levels. One possibility is that this is due to mechanical cues: it has been shown that spatial confinement of cells in a three-dimensional microenvironment results in the maintenance of pluripotency even in the absence of LIF ^48^. In the early mouse embryo, we might be observing a similar effect. Another possibility is that the Hippo signalling pathway is involved in interpreting the positional information as previously suggested ^49^. Hippo signalling is clearly determining the first fate choice (trophectoderm versus ICM) in the mouse embryos ^50–54^ and it has been shown that the second fate choice (Epi versus PrE) is linked to the first one ^55^. Furthermore, Hippo signalling regulates NANOG expression in mouse embryonic stem cells ^56^. Our results are consistent with mechanical cues as well as Hippo signalling being the driver of expression of higher levels of NANOG in the central cells and their differentiation into Epi cells. Furthermore, we observed that in the absence of GATA6, the spatial distribution of NANOG expressing cells is disrupted. These results show that besides the known inhibition of NANOG by GATA6 within a cell ^7, 57, 58^, GATA6 is also involved in regulating NANOG expression in the cell neighbourhood.

The spatial pattern of cells depending on their GATA6 expression levels correlates with a local neighbourhood. In early embryos, cells with similar GATA6 expression levels tend to be found close to one another (and this distribution is independent of their localization within the ICM). The functional relevance of the clustering effect might be to ensure a coordinated PrE cell behaviour during their migration to occupy their final position at the blastocoele. Since GATA6 expression in this stage is regulated through FGF/MAPK signalling via FGFR1 ^14, 15^, this pathway might be involved also in regulating the spatial distribution of GATA6 expressing cells. Furthermore, differential activity of FGF/MAPK via FGFR1 might also be involved in regulating the spatial three-dimensional patterns of NANOG expressing cells in early blastocysts ^14, 15^. It has been shown that a subpopulation of ICM cells produce FGF4 in this stage ^13^. Based on our results, we hypothesise that the secretion of FGF4 starts in the subpopulation of NANOG positive cells with 9 neighbours and positioned away from the centroid. Binding of FGF4 to FGFR1 will downregulate NANOG and promote GATA6 expression.

In summary, there must be a coordinating mechanism regulating NANOG and GATA6 expressing cells three-dimensional distribution in early blastocysts. The primary signalling pathway involved is probably FGF/MAPK signalling via FGFR1 ^14, 15^. However, we cannot rule out that other major signalling pathways involved in patterning fields or groups of cells, such as Notch, Wnt, BMP, or EGF might also have an input ^59^. In support of this, there are reports of Notch signalling involved in early mouse development ^60–62^, as well as Wnt signalling ^63^, BMP signalling ^64, 65^, EGF signalling ^66^. In the light of our results, it will be important to revisit how these signalling pathways might be involved in cell fate decision in early blastocysts investigating how they affect cell position within the ICM.

### Spatial pattern of NANOG and GATA6 expressing cells in the ICM of mid/late blastocysts

As development progresses, we observed that the three-dimensional distribution of ICM changes. In mid blastocysts, the highest NANOG expressing cells still have around 9 neighbours, but now they occupy central locations within the ICM (Fig. 8). In our results, we cannot distinguish whether central cells upregulate NANOG or high NANOG cells actively move to the centre of the ICM. However, previous studies using a fluorescent reporter line showed active migration of PrE-to-Epi converted cells towards inner regions of the ICM, supporting an active migration ^27^. In this stage, there might still be some input from mechanical cues and/or Hippo signalling regulating NANOG expression (see above). Regardless of whether a passive or active mechanism is involved, the central location of high NANOG cells implies that those cells will acquire Epi fate by more efficiently inhibiting GATA6 expression, maintaining their position in late blastocysts ^4, 5, 7–11^. The analysis of the position of GATA6 expressing cells within the ICM in mid blastocysts reveals that they are already localised next to the blastocoele (Fig. 8). This is consistent with previous studies showing that PrE progenitors tend to maintain their original position ^3^.

Previous studies suggested that NANOG and GATA6 are randomly or alternatingly distributed in the ICM of mid blastocysts ^2^. Our results show that both NANOG positive and negative cells as well as GATA6 positive and negative are not distributed randomly or follow a clear alternating pattern. Only the analysis with the first data set showed that GATA6 positive and negative cells followed a combination of the random and period two patterns. These results explain why, if a small data set is used to interrogate the system, it could lead to the proposal of a random or alternating model for the distribution of NANOG and GATA6 positive cells during Epi versus PrE fate differentiation.

Epi progenitors secrete FGF4 which binds to FGFR2 expressed in PrE progenitors reinforcing their decision ^12–15^. Here we could rule out that Epi progenitors instruct PrE fate in the direct neighbours. This poses the question of how FGF4 is propagated extracellularly once secreted in preimplantation embryos which prevents its activity in the direct neighbours. One possibility is that it is via differential expression of heparan sulfate (HS) chains which has been associated with heterogeneous di-phosphorylated Erk (FGF signalling activity readout) in this embryonic stage ^67^. Another alternative is changes in the internalization and spreading related to endocytosis rates as shown for FGF8 in zebrafish embryos ^68^. These possibilities are not the only ones and just represent ways in which FGF4 might reach Epi and PrE progenitors differently eliciting different responses.

In summary, our work documents the first analysis of the three-dimensional spatial distribution of NANOG and GATA6 expression in the ICM of the mouse blastocyst. Traditionally, cell fate decision has been studied in the context of signalling pathways and how these regulate fate markers expression. Here, we include a new aspect, namely the cell neighbourhood and how it is varies depending on the cell fate. This cell neighbourhood is characterised by the expression levels of the fate markers in the surrounding cells, together with the number of surrounding cells and cell position. Our results establish the importance of studying intercellular interactions and highlights that Epi versus PrE differentiation is initialised in early blastocysts. In the future, besides studying how other signalling pathways might be involved in the regulation of the cell neighbourhood, it would be interesting to simulate current theoretical models of Epi versus PrE differentiation behave when simulating them in three-dimensional cell clusters.

## Materials and Methods

### Ethics statement

All mouse work was approved by the University of Bath Animal Welfare and Ethical Review Body (AWERB) and undertaken under UK Home Office license PPL 30/3219 in accordance with the Animals (Scientific Procedures) Act incorporating EU Directive 2010/63/EU.

### Mice, embryos and immunohistochemistry

Wild-type CD1 embryos were generated by in-house breeding and natural mating. Detection of copulation plug confirmed successful mating; the resulting embryos were then considered to be embryonic day (E) 0.5. Embryos were isolated in M2 medium (Millipore). Embryos were prepared for immunofluorescence as previously described (Nichols et al., 2009). Primary antibodies used were: anti-NANOG (eBiosciences 14-5761, 1:100), anti-GATA6 (R&D, AF1700, 1:200). Nuclei were stained using DAPI or Hoechst (1:500, Invitrogen). Embryos were mounted on microscopy slides with Vaseline bridges to prevent their crushing. Three independent immunofluorescence stainings, each with E3.5 and E4.5 embryos from 7 litters, were performed.

### Imaging and automated image analysis

A total of 45 embryos was imaged in four batches of 19, 15, 2 and 9 embryos. Images were acquired using a Zeiss LSM 510-META and a Plan-Apochromat 63x/1.4 Oil Ph3, with optical section thickness of 1 μm. All images in each imaging session were obtained using the sequential scanning mode, with the same conditions of laser intensity, gain, and pinhole, and were processed in exactly the same way. The range indicator palette option (Zeiss AIM software) was used to ensure that no oversaturated images were taken. The three-dimensional image stacks were segmented using MINS ^30^, cells were semi-automatically assigned to ICM or TE, the features of the cell nuclei were extracted including the nuclear centroid and volume, together with the mean intensity of NANOG and GATA6 for each nucleus.

### Automated image pre-processing, normalization and classification of cell populations

We checked for fluorescence intensity decay along the z-axis. As this decay was minimal due to the mounting of the embryos, intensity adjustment along z was not performed (not shown).

The mounting of the embryos resulted in a slight squeezing along the z-axis of the image and hence extension along x and y. We assumed that the embryos that do not have fully segregated epiblast and primitive endoderm should be spherical (early and mid blastocysts, see below). Based on this assumption, we calculated the deviation from sphericity for each of these embryos and rescaled the coordinates of the cell nuclei to obtain spherical embryos. Embryos with segregated epiblast and primitive endoderm have hatched from the zona and are elongated (late blastocysts, see below). To rescale the coordinates of these embryos, we calculated for each batch the median rescaling factors for x, y and z of the early and mid blastocysts and applied these to the late ones.

Plotting the mean GATA6 expression levels versus the mean NANOG expression levels for all nuclei, we observed a shift in the data related to the batch number, each batch corresponding to a different imaging session (Fig. 2A). To align the data obtained from the four independent sessions, we focused on the embryos with fully segregated Epi and PrE (late blastocysts). We performed a manually curated k-means clustering of the data and obtained thresholds for NANOG and GATA6 for each batch. Based on these thresholds we calculated transformations to align the intensity values of all data sets (all batches and all stages). This changes the absolute intensity levels for each embryo but it does not change the relative intensity values in an embryo which is the value that is relevant for our analysis. Based on the thresholds for GATA6 and NANOG, all cells were classified as double negative (DN, N- and G-), NANOG+/GATA6-(N+/G-), NANOG-/GATA6+ (N-/G+) and double positive (DP, N+, G6+). The calculations were performed with Mathematica 11.1 (Wolfram Research).

### Embryo staging

We separated the embryos from our experiments into three developmental stages. Embryos with clearly separated Epi and PrE were set to be late blastocyst stage. To separate the remaining embryos in early and mid blastocyst stage, we applied staging by cellular event. This staging method is based on the progression of the embryos in terms of Epi versus PrE differentiation. Therefore, the developmental stage of the embryo is determined based on the percentage of cells expressing both NANOG and GATA6 (DP). Embryos with the majority, hence more than 55%, DP cells in the ICM were classified as early blastocyst stage and embryos with more than 0% but less than 55% DP cells were classified as mid blastocyst stage. The calculations were performed with Mathematica.

### Pre-processing and staging of data from Saiz *et al.* 2016 ^24^

We used the embryos labelled as “littermate”, available from GitHub ^24^. This resulted in 147 additional data sets. To identify early, mid and late blastocysts we applied the staging by cellular event as indicated above.

From these data, we excluded all NANOG and GATA6 levels from the distribution that were two standard deviations away from the respective mean as we noticed that there were some oversaturated nuclei images. The remaining analysis was analogous to our data. The calculations were performed with Mathematica.

### Cell graph generation and neighbourhood analysis

We derived a cell graph representation to characterise the spatial distribution of the cells in each embryo in our data set, the data set from ^24^ and the data set from ^6^ composed by 20 *Gata6^+/+^* embryos, 28 *Gata6+/-* embryos, and 15 *Gata6^-/-^* embryos. Delaunay cell graphs were generated ^29^ based on the pre-processed nuclei coordinates. The Delaunay cell graph (DCG) is given by *DCG*(*V*, *E_DCG_*) where *V* is the vertex set and *E_DCG_* is the edge set of the graph. The DCG graph is constructed based on a Delaunay triangulation. Delaunay triangulation and its dual, the Voronoi tessellation are routinely used to approximate which cells are in physical contact ^29, 31, 32^. An edge (*u*, *w*) was created between two vertices *u* and *w* if the corresponding points are connected by a line in the Delaunay triangulation and the Euclidean distance between *u* and *w* was less than 30μm.

For a given cell (vertex in the cell graph), the neighbouring cells are represented by the vertices that are connected through edges. We exploited a positive or negative cell state for ICM cells and their neighbours within the ICM for Figure 3C-D, and Figures S2, S4, and S8A-F. For Figures 4, 5, 6 and 7 as well as Figures S5, S7, S8G-H and S11A-D, we include all neighbours, i.e. in this case TE cells can occur as neighbours. The calculations were performed with Mathematica.

### Simulations of salt-and-pepper patterns

We generated three simulated patterns of expression level distribution, a random expression pattern, a period two pattern and a nearest neighbour pattern. For the simulations of the patterns, we used the positions of the cells in the ICM of embryos at early and mid blastocyst stage. Hence, for each blastocyst, we obtained three simulated expression patterns.

For the random expression pattern of N+ and N-cells, we extracted the distribution of NANOG levels for the ICM cells of all embryos. Based on this distribution, we randomly assigned simulated expression levels to the cells of the ICM of early and mid blastocyst embryos. Then, we separated the cells with the random expression levels into positive and negative cells based on the thresholds for NANOG obtained during the pre-processing of the data. Finally, we used the cell graph to determine how many negative neighbours in the ICM a positive cell has and vice versa. Similarly, we obtained the distribution of GATA6 levels of ICM cells in all embryos, randomly assigned expression levels to the ICM cells of early and mid blastocyst embryos, separated the cells in G6+ and G6- and calculated the distributions of positive and negative neighbours in the ICM. Using the actual data as a basis ensured that we really only test the randomness of the distribution of cell fates and do not get any bias from potential assumptions for the NANOG and GATA6 distributions that we would need to make if we were using theoretical distributions for NANOG and GATA6.

For the period two pattern, we generated for each ICM a pattern in which cells in a positive state have only cell neighbours in a negative state. For the nearest neighbour pattern, we started with one cell and assigned it a negative state. Then, we chose the nearest neighbour cell based on the Euclidean distance between the two nuclei centroids that did not have a state yet and assigned it a positive state. We iterated this procedure until all cells were assigned a state. The simulations were performed with Mathematica.

### Correlation analyses

To relate the expression levels of a given cell to the expression levels of all its neighbours (both TE and ICM), we calculated Spearman’s correlation coefficient of the expression levels of a cell and the median expression levels of its neighbours. The Spearman correlation coefficient was used, because the pairs of analysed variables are not normally distributed (not shown, ^47^).

To determine whether the obtained correlations are statistically significant, we performed a bootstrap resampling of the correlation coefficients of our data and compared the result with correlation coefficients of a random model. For the bootstrapping, we resampled the experimental data to create 10,000 different data sets. The random expression model was calculated for NANOG and GATA6 in all embryos. Based on the distribution of NANOG expression levels of ICM cells of all embryos, we randomly assigned simulated expression levels to all cells of early, mid and late blastocyst embryos. Then, we separated the ICM cells with the random expression levels into the four populations DN, DP, N+/G6-, N-/G6+ based on the thresholds for NANOG obtained during the pre-processing of the data. Finally, we used the cell graph to determine the neighbours of a given cell and their expression levels. For GATA6 we took a similar approach. The Spearman’s correlation analysis and bootstrapping were performed in Matlab R2012b. The simulations of the random expression model were performed with Mathematica.

To classify the strength of the correlations we used the criteria by Evans ^69^:

0.00-0.19: ‘very weak’

0.20-0.39: ‘weak’

0.40-0.59: ‘moderate’

0.60-0.79: ‘strong’

0.80-1.0: ‘very strong’.

### Statistics

Mann-Whitney tests or z-tests with Bonferroni corrections were performed in Matlab ^47^.

## Acknowledgements

Research in the AK Stelzer is supported by the Deutsche Forschungsgemeinschaft (DFG, CEF-MC II, EXC-115).). Research in the Muñoz-Descalzo lab is supported by the University of Bath and a Wellcome Trust Seed Award (109589/Z/15/Z). SCF and SMD acknowledge the support by an International Exchanges Grant from The Royal Society (IE141022). We thank Christian Schröter, Jennifer Nichols, Joaquín de Navascués and Alfonso Martínez-Arias for helpful comments. We also want to that Néstor Sáiz and Kat Hadjantonakis for their comments, sharing their quantitative data through GitHub as well as through personal communication. The funders had no role in study design, data collection and analysis, decision to publish, or preparation of the manuscript.

## Supplementary Figure Legends

**Fig. S1.**
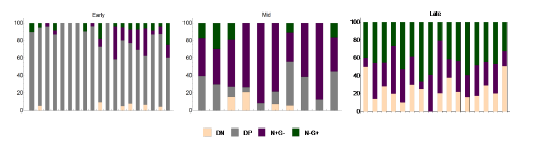
Population analyses in ICMs of individual embryos staged by event. Early, mid and late blastocysts population distributions in ICM in single embryos staged by proportion of DP cells.

**Fig. S2.**
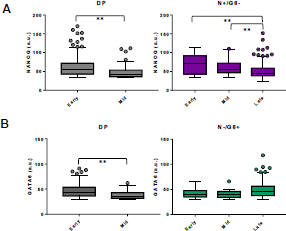
Expression levels for each population in ICM cells for both staging methods. **(A-B)** Box plots showing the expression levels of NANOG (A) and GATA6 (B) in the indicated stages and populations. Mann-Whitney test with Bonferroni correction between the different populations; **:p<0.01.

**Fig. S3.**
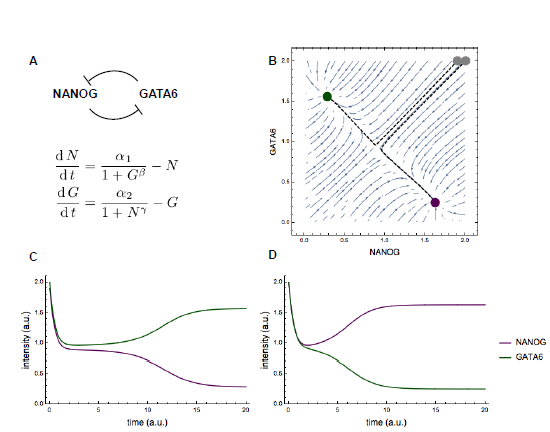
Illustration of a bistable switch behaviour. **(A)** System of equations for a bistable switch for NANOG expression (N) and GATA6 expression (G). **(B)** Exemplary phase diagram for the system in (A) for parameter values α1=1.64, α2=1.585, β=3.5,γ=3.46. The dashed black lines indicate the trajectories for the solutions for the initial values [N,G]=[1.9,2] (steady state in green) and [N,G]=[2,2] (steady state in purple). **(C)** Plot of the levels of NANOG (purple) and GATA6 (magenta, y-axis) over time (x-axis) for an initial NANOG value of 1.9 and an initial GATA6 value of 2. **(D)** Plot of the levels of NANOG (purple) and GATA6 (magenta, y-axis) over time (x-axis) for initial NANOG and GATA6 values of 2.

**Fig. S4.**
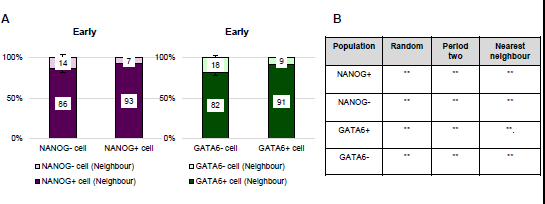
Spatial analysis of cell fate markers in early embryos. **(A)** Proportion of N+ or N-neighbours of N+ or N-cells (left) and G6+ or G6-neighbours of G6+ or G6-cells (right) in the ICM of early blastocyst embryos. Error bars correspond to the standard error of the mean. **(B)** Table summarizing the results of the Mann-Whitney statistical tests with Bonferroni correction comparing the simulated distributions and experimental data obtained in the early blastocysts. **: p<0.01.

**Fig. S5.**
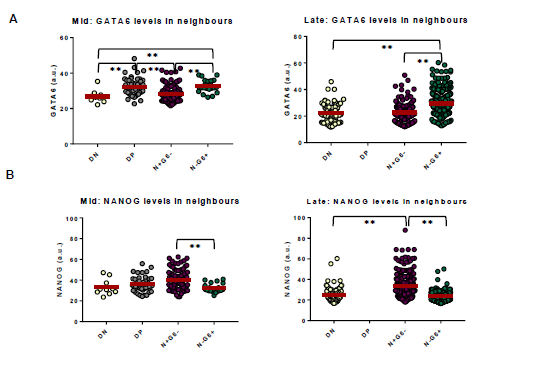
NANOG and GATA6 expression levels in neighbouring cells. **(A)** Scatter dot plots showing the expression levels of GATA6 in neighbours of cells for the four populations in early, mid and late blastocysts. Mann-Whitney test with Bonferroni correction between the different populations: **: p<0.01. **(B)** Scatter dot plots showing the expression levels of NANOG in neighbours of cells for the four populations in early, mid and late blastocysts. Mann-Whitney test with Bonferroni correction between the different populations: **: p<0.01.

**Fig. S6:**
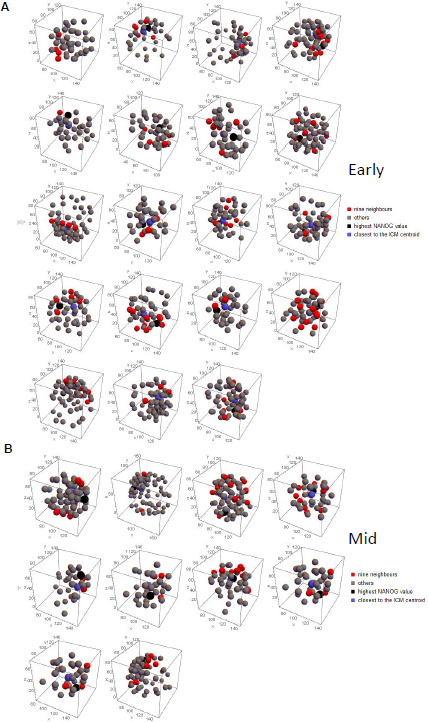
Relative cell positions in individual early and mid blastocysts. (**A-B**) Three-dimensional representation of each early (A) and mid (B) blastocyst from the data set. Cells with 9 neighbours are shown in red, cell closest to the ICM centroid is in blue, the cell expressing the highest NANOG levels is shown in black, and other cells in grey. **(B)** Table summarizing the results of the Mann-Whitney tests with Bonferroni correction comparing the simulated distributions for the three prevailing notions of salt-and-pepper pattern and the experimental data obtained in the mid blastocysts. **: p<0.001. See also Fig. S7A-C.

**Fig. S7.**
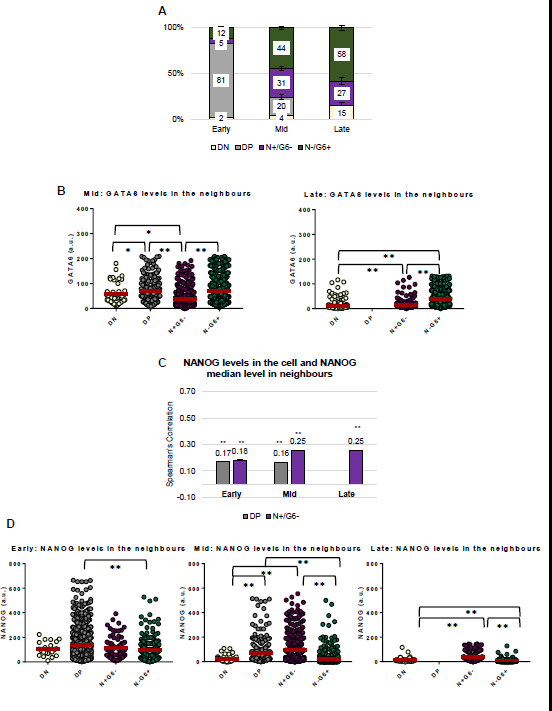
Population, expression levels and NANOG correlation analyses from ^24^ data set. **(A)** Proportion of cells expressing GATA6 (N-/G6+), NANOG (N+/G6-), co-expressing NANOG and GATA6 (double positive, DP) or neither (double negative, DN) in ICM cells at different developmental stages. Staging was performed according to the percentage of double positive cells as shown in Fig. 2C. **(B)** Scatter dot plots showing the expression levels of GATA6 in the indicated populations in mid (left) and late (right) blastocysts. Mann-Whitney test with Bonferroni correction between the different populations: **: p<0.01; *: p<0.05. **(C)** Spearman’s correlation coefficients of NANOG levels of a cell and the median NANOG levels of its neighbours in the indicated populations in the ICM at different embryonic developmental stages. **: p<0.01, Mann-Whitney test with Bonferroni corrections. **(D)** Scatter dot plots showing the expression levels of NANOG (B) in neighbours of cells for the four populations in early (left), mid (centre) and late (right) blastocysts. Mann-Whitney test with Bonferroni correction between the different populations: **: p<0.01; *: p<0.05.

**Fig. S8:**
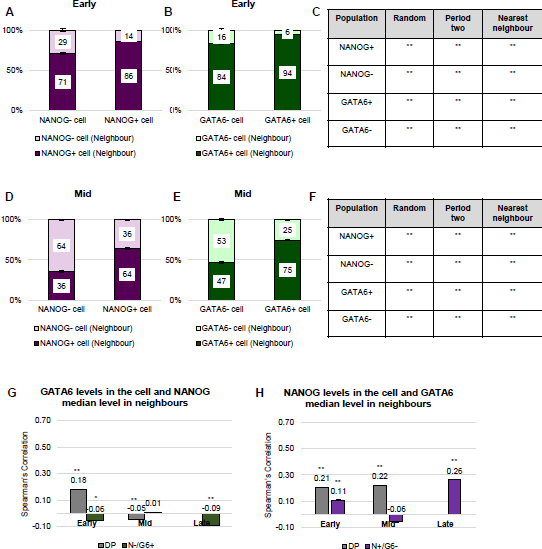
Distribution of positive and negative cells in early and mid blastocysts and extended correlation analyses from ^24^ data set. (**A-B**,**D-E**) Measurement of percentage of N+ or N-neighbours of N+ or N-(A,D) and G+ or G6-neighbours of G6+ or G6-(B,E) in the ICM of early (A,B) and mid blastocysts (D,E). **(C,F)** Summary results of the Mann-Whitney statistical tests with Bonferroni correction comparing the simulated and experimental data proportions obtained in the early (C) and mid (F) blastocysts. **:p<0.01. **(G-H)** Spearman’s correlation coefficients of GATA6 levels of a cell and the median NANOG levels of its neighbours (G) and NANOG levels of a cell and the median GATA6 levels of its neighbours (H) in the indicated populations in the ICM at different embryonic developmental stages. **: p<0.01; *: p<0.05, Mann-Whitney test with Bonferroni corrections.

**Fig. S9:**
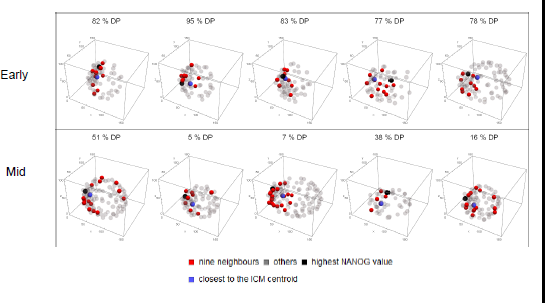
Relative cell positions of selected individual early and mid blastocysts from ^24^ data set. Three-dimensional representation of 5 representative early and mid blastocyst from the data set. The percentage of DP cells in each embryo is shown. Cells with 9 neighbours are shown in red, cell closest to the ICM centroid is in blue, the cell expressing the highest NANOG levels is shown in black, and other cells in grey.

**Fig. S10:**
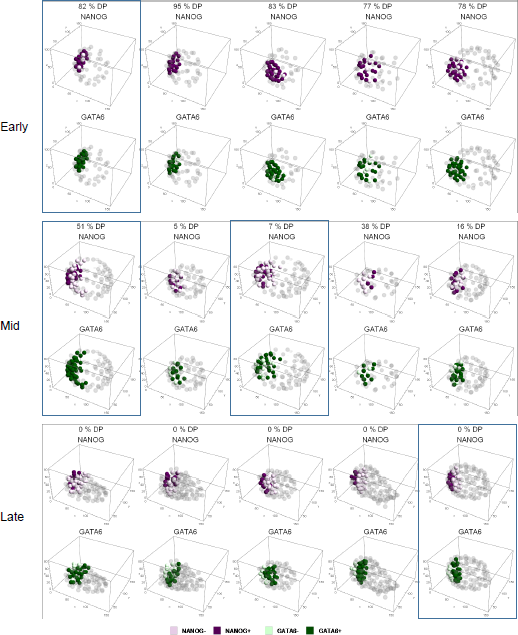
Relative cell positions of N+/N- and G6+/G6-in selected individual early, mid and late blastocysts from ^24^ data set. Three-dimensional representation of 5 representative early, mid and late blastocyst from the data set. The embryos in the blue boxes show the cell distribution represented in Fig. 8. The percentage of DP cells in each embryo is shown. N+ are in dark magenta; N-, in pink; G6+, in dark green, G6-, in light green; TE, in grey.

**Fig. S11:**
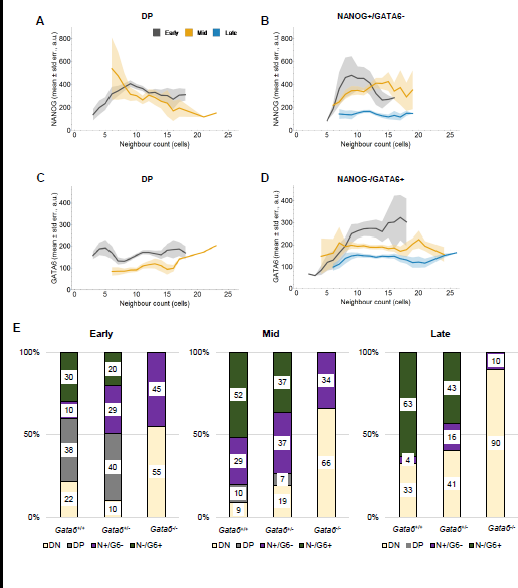
Relationship between NANOG or GATA6 expression levels and number of neighbours in different populations and stages and population analysis from ^6^ data set. **(A-B)** Mean level of NANOG (y-axis) versus the number of neighbours (x-axis) for ICM cells in DP (A) and N+/G6-(B) cells. **(C-D)** Mean level of GATA6 (y-axis) versus the number of neighbours (x-axis) for ICM cells in DP (B) and N-/G6+ (B’) cells. The shaded regions display the standard error of the mean. **(A)** Proportion of cells expressing GATA6 (N-/G6+), NANOG (N+/G6-), co-expressing NANOG and GATA6 (double positive, DP) or neither (double negative, DN) in ICM cells at different developmental stages in *Gata6^+/+^, Gata6^+/-^* and *Gata6^-/-^* Staging was performed as in ^6^.

